# Learning brain dynamics for decoding and predicting individual differences

**DOI:** 10.1101/2021.03.27.437315

**Authors:** Luiz Pessoa, Chirag Limbachia, Joyneel Misra, Srinivas Govinda Surampudi, Manasij Venkatesh, Joseph Jaja

**Affiliations:** Department of Psychology and Maryland Neuroimaging Center, University of Maryland, College Park, MD, USA; Department of Electrical and Computer Engineering, University of Maryland, College Park, MD, USA

## Abstract

Insights from functional Magnetic Resonance Imaging (fMRI), as well as recordings of large numbers of neurons, reveal that many cognitive, emotional, and motor functions depend on the multivariate interactions of brain signals. To *decode* brain dynamics, we propose an architecture based on recurrent neural networks to uncover distributed spatiotemporal signatures. We demonstrate the potential of the approach using human fMRI data during movie-watching data and a continuous experimental paradigm. The model was able to learn spatiotemporal patterns that supported 15-way movie-clip classification (~90%) at the level of brain regions, and binary classification of experimental conditions (~60%) at the level of voxels. The model was also able to learn individual differences in measures of fluid intelligence and verbal IQ at levels comparable or better than existing techniques. We propose a dimensionality reduction approach that uncovers low-dimensional trajectories and captures essential informational (i.e., classification related) properties of brain dynamics. Finally, saliency maps were employed to characterize brain-region/voxel importance, and uncovered how dynamic but consistent changes in fMRI activation influenced decoding performance. When applied at the level of voxels, our framework implements a dynamic version of multivariate pattern analysis. We believe our approach provides a powerful framework for visualizing, analyzing, and discovering dynamic spatially distributed brain representations during naturalistic conditions.^1^

**Author summary:** Brain signals are inherently dynamic and evolve in both space and time as a function of cognitive or emotional task condition or mental state. To characterize brain dynamics, we employed an architecture based on recurrent neural networks, and applied it to functional magnetic resonance imaging data from humans watching movies or during continuous experimental conditions. The model learned spatiotemporal patterns that allowed it to correctly classify which clip a participant was watching based entirely on data from other participants; the model also learned a binary classification of experimental conditions at the level of voxels. We developed a dimensionality reduction approach that uncovered low-dimensional “trajectories” and captured essential information properties of brain dynamics. When applied at the level of voxels, our framework implements a dynamic version of multivariate pattern analysis. We believe our approach provides a powerful framework for visualizing, analyzing, and discovering dynamic spatially distributed brain representations during naturalistic conditions.

## Introduction

As brain data become increasingly *spatiotemporal*, there is a great need to develop methods that can effectively capture how information across space and time support behavior. In the context of functional Magnetic Resonance Imaging (fMRI), although data are acquired temporally, they are frequently treated in a relatively static manner (e.g., in event-related designs, responses to short trials are estimated with multiple regression assuming canonical hemodynamics). However, a fuller understanding of the mechanisms that support mental functions necessitates the characterization of *dynamic* properties, particularly during the investigation of more naturalistic paradigms, including movie watching [1], and other continuous paradigms [2, 3].

A central goal of neuroscience is to *decode* brain activity: to infer mental processes and processes from brain signals. In fMRI, most approaches are spatially constrained and consider voxels in a local neighborhood (”searchlight”) or across a few regions. Such models are useful to decode stimuli or mental state, particularly when the problem is well understood and localized anatomically [4–7]. Additional techniques decode brain activity based on whole-brain functional connectivity matrices [8], as well as machine learning methods [9]. The vast majority of decoding methods are static, that is, the inputs to classification are patterns of activation that are averaged across time (“snapshots”) [10]. Some studies have proposed using temporal information in addition to spatial data [11–16]. In such cases, features used for classification are extended by considering a temporal data segment instead of, for example, the average signal during the acquisition period of interest.

Despite some progress in utilizing temporal information in decoding methods, key issues deserve to be further investigated:

- How can we characterize brain signals generated by dynamic stimuli in terms of generalizable (across participants) spatiotemporal patterns? How are such patterns distributed across both space and time? For example, although a recent study used recurrent neural networks to decode brain states from fMRI data, they investigated different working memory conditions (remembering faces, bodies, tools, or places) which are associated with stable brain states at the temporal scale of technique [17].
- Understanding the dimensionality of brain representations has become an important research question in recent years [18–20]. For example, for simple movements, neuronal signals across populations of neurons can be well approximated by a projection onto a two-dimensional representation [21]. This question is now starting to be addressed in the fMRI literature [22, 23]. Here, we tackle the problem of learning low-dimensional spatiotemporal signals from dynamic stimuli, including movie watching and continuous experimental paradigms.
- If spatiotemporal patterns capture important properties of brain dynamics, do they capture information about individual differences that are predictive of behavioral capabilities?
- Although multivariate techniques, including neural networks, can be successful at the level of prediction, *interpretability* is of key importance in the context of neuroscience. Therefore, here we quantified the relative importance of spatiotemporal features, specifically how brain regions contribute to input classification as a function of time.
- Can spatiotemporal decoding be applied at both the region-of-interest and voxel levels? Extending it to the voxel level allows investigators to probe finer spatial representations underlying task conditions or states of interest, and opens the way for dynamic representational similarity analysis [24].

In the present paper, we sought to address the above challenges by developing a *dynamic* computational framework built upon recurrent artificial neural networks.

## Materials and methods

### Data

#### Movie data

We employed Human Connectome Project (HCP; [25]) data of participants scanned while watching movie excerpts (Hollywood and independent films) and other short videos, which we call “clips”. The legend of Fig 4 provides further information. All 15 clips contained some degree of social content. Participants viewed each clip once, except for the *test-retest* clip that was viewed 4 times. Clip lengths varied from 65 to 260 seconds (average: 198 seconds).

We employed all available movie-watching data, except those of 8 participants with runs missing; thus we used *N* = 176 participants. Data were sampled every 1 second. The preprocessed HCP data included FIX-denoising, motion correction, and surface registration [26, 27]. We analyzed data at the region of interest (ROI) level, with one time series per ROI (average time series across locations). We employed a standard 300-ROI cortical parcellation [28]. Thus, the input time series consisted of a vector of *N_x_* = 300 dimensions at each time, *t*. Individual ROI time courses were normalized by z-scoring (i.e. centered to zero mean and unit standard deviation) to help remove potential differences across runs/participants.

#### Moving-circles paradigm

We employed data from a separate dataset collected in our laboratory [3]. Briefly, the experimental paradigm was as follows. Over a period of 3 minutes, two circles moved on the screen, at times approaching and at times retreating from each other. If they touched, participants received a non-painful, but highly unpleasant electrical shock. The movement of the circles was smooth but not predictable. In particular, the circle motions were set up with multiple instances of what we called “near misses”: periods of approach followed by retreat; in such instances, the circles came close to each other and retreated just before colliding. Based on near-misses, we defined *approach* and *retreat* states: each lasted 6-9 seconds during which the circles approached or retreated from each other.

Data from *N* = 61 participants were employed, and were sampled every 1.25 seconds. Preprocessing steps included motion correction (ICA-AROMA via FSL). We analyzed data at the voxel level from the left anterior insula, as defined according to an anatomical template [29]. The input time series consisted of a vector of *N_x_* = 745 dimensions at each time, *t*.

### Performance

#### Accuracy

At each point in time, *t*, we computed accuracy as the average true positive rate (TPR) across inputs of a given class (movie clip or experimental condition). Specifically, for each class label *c*:

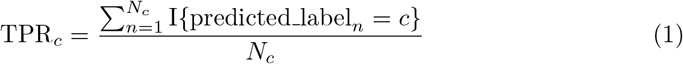

where *N_c_* is the total number of data samples belonging to class *c*, predicted label_*n*_ is the label predicted by the model for the *n*^th^ data sample, and I is an indicator function which equates to 1 if predicted label_*n*_ = *c* and 0 otherwise. We computed accuracy at each time step and visualized it as a function of time (see Fig 3).

#### Training/Testing split and Cross-Validation

Movie data from 100 participants were used for training, and the remaining 76 participants were used as a test set (thus completely invisible to model tuning/learning). To determine optimal hyperparameters for clip prediction, we employed a 5-fold cross-validation approach on the training set. In each fold, participants in the training set were not included in the validation set. After determining optimal hyperparameters, we retrained the model on the entire training data. Final results were generated exclusively based on test data. A similar approach was used for the the moving-circles paradigm, where data from 42 and 19 participants were used as a training set and test set, respectively. For cross validation, the folds had size {9, 9, 8, 8, 8}.

### Basic recurrent architecture for classification

Recurrent neural networks allow signals at the current time step to be influenced by long-term past information. Although it was originally difficult to develop effective learning procedures for these algorithms (e.g., vanishing/exploding gradients prevented them from learning relationships beyond 5-10 time steps; [30]), recent developments have largely overcome these challenges. Here, we employed a recurrent neural network architecture based on Gated Recurrent Units (GRUs; [31]). Fig 1A shows the model. Briefly, inputs ***x**_t_* were provided to a GRU network that produced a “hidden” representation, ***h**_t_*, of the incoming signal based on current and past signals. For classification purposes, the hidden-layer signals were transformed to create a vector of class scores, the maximum of which determined the model’s prediction of the input. Next, we describe the model formally.

**Fig 1.**
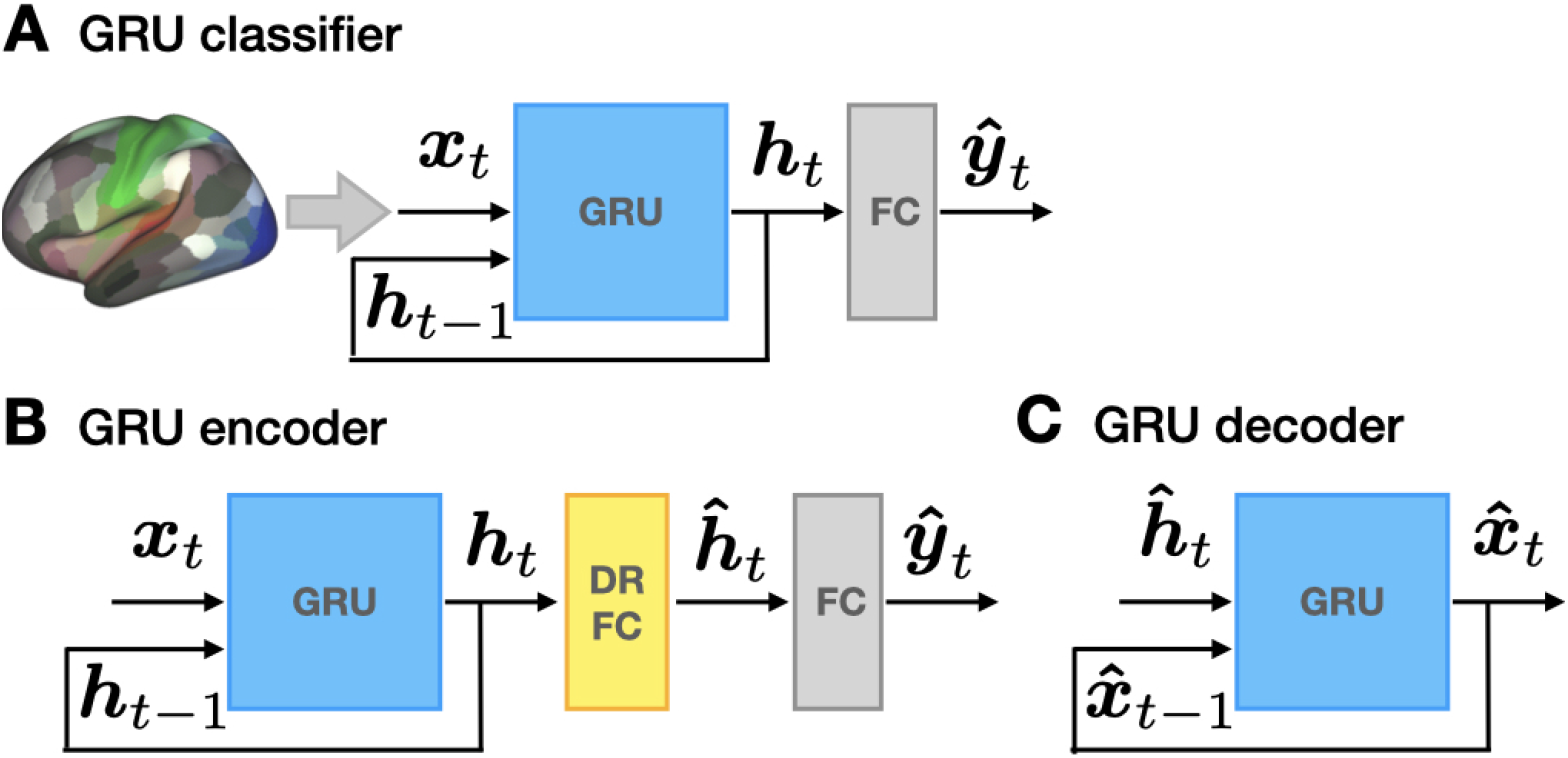
Model architectures based on neural networks with gated recurrent units (GRUs). (A) *Classifier*. At each time step, time series data,***x**_t_*, provided inputs. The recurrent neural network transformed the inputs into a latent representation, ***h**_t_*, which then determined the output class scores, ***ŷ**_t_*. The unit with highest activation determined the model’s prediction of the input stimulus at each time. (B) *Dimensionality reduction*. GRU outputs, ***h**_t_*, were first linearly projected to a lower-dimensional space using a fully-connected layer (DR-FC). Classification was then performed based on the low-dimensional representation, ***ĥ**_t_*. (C) *Input signal reconstruction*. A separate GRU was trained independently to reconstruct the original brain signals based on the low-dimensional signals, ***ĥ**_t_*.

Time series data from units of interest (ROIs or voxels) comprised the input vector at each time, ***x**_t_*. The GRU network (with one or more layers) transformed its input, ***x**_t_*, non-linearly onto its output, ***h**_t_*, the hidden layer of the classifier system (see S1 Appendix for details). Because our goal was input classification, GRU outputs at each time, ***h**_t_*, passed through an additionally fully-connected (FC) layer of units generating a vector of class scores:

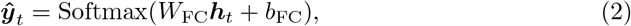

where Softmax is the generalized logistic activation function. As the outputs are in the range [0, 1] and and add up to 1, they can be treated as probabilities. The output of the model was the unit label, *i*, such that it selected the class with largest probability: 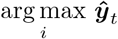. As all operations were performed at every time, *t*, the final output was a time series of label predictions. For movie clips, we employed data from 300 ROIs, so the dimensionality of the input was *N_x_* = 300. The GRU network employed a single layer with 32 units (*N_h_* = 32). The dimensionality of ***ŷ**_t_* was *N_y_* = 15 to implement 15-way classification (based on the number of clips). For the moving-circles data, we employed data from 745 voxels (*N_x_* = 745). The GRU network contained three layers of 32 nodes each; the third layer corresponded to the vector ***h**_t_* (*N_y_* = 32). The dimensionality of ***ŷ**_t_* was *N_y_* = 2 to implement threat vs. safe classification.

All weight matrices, *W*, required for GRU tuning (see Appendix 1) and the bias parameters, *b*, underwent supervised training. Training sought to minimize the cross-entropy (CE) loss between predicted labels, ***ŷ**_t_*, and true labels, ***y**_t_* (as provided by the supervisor), and was evaluated at at each time step:

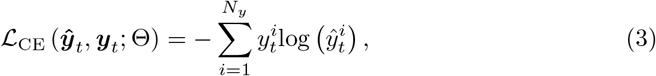

where the superscript *i* indexes the dimension of the predicted and teaching signals. The set of trainable parameters was denoted as Θ. The cross-entropy attains its minimum value when the two distributions, ***ŷ**_t_* and ***y**_t_*, are identical. We defined the total loss function, *J* (Θ), as the average cross-entropy loss across time points (i.e., the average value across time of Eq 3), thus encouraging predicted labels to be as close to the true labels as possible across time. We minimized *J* (Θ) using the backpropagation through time algorithm [32]; gradient descent was optimised with the Adam optimizer [33]. The model was implemented using TensorFlow [34].

As our goal was not necessarily maximizing model performance, we explored a small set of potential architectures by varying the number of hidden layers and number of units per layer (Fig 2). For the region-based analysis, we explored the {1, 2, 3} × {16, 32, 64} space (units by layers). As accuracy was relatively high overall (*>*70%) and improvements were relatively modest as a function of these parameters, we chose the simplest case (one layer) but with 32 units which improved performance over 16 units (for evaluation performance, we considered all time points, even those during the first 60-90 seconds when accuracy increased sharply; see Fig 3). For the voxel-based analysis, we explored the {1, 2, 3} × {16, 32, 64, 128, 256} space. Classification of approach vs. retreat was a challenging problem, so we utilized 3 layers; we employed 32 units per layer for consistency with the clip classification architecture.

**Fig 2.**
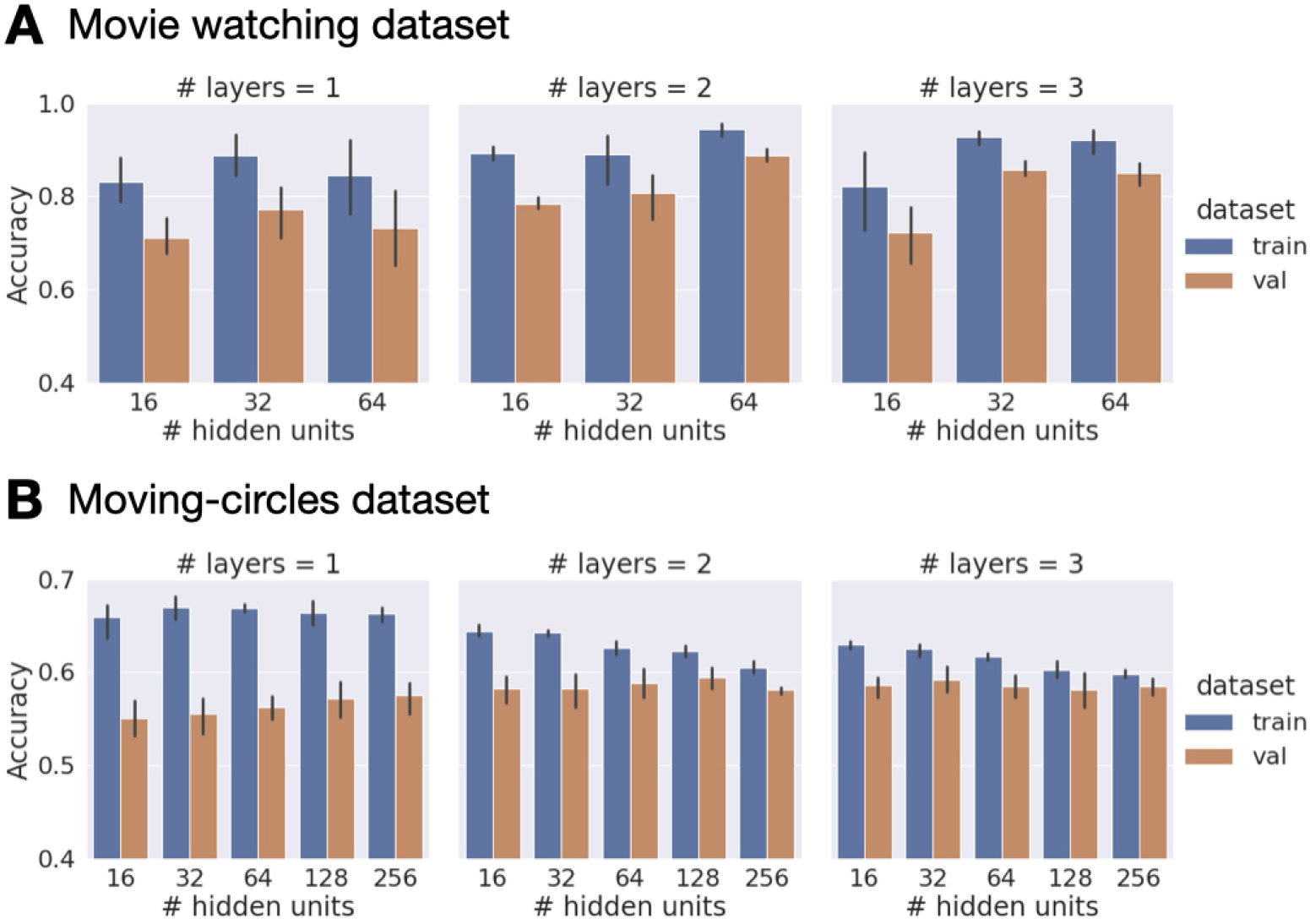
Selection of model architectures via cross validation on data from 100 participants. Val: validation portion of training data. Error bars show 95% confidence intervals across folds.

**Fig 3.**
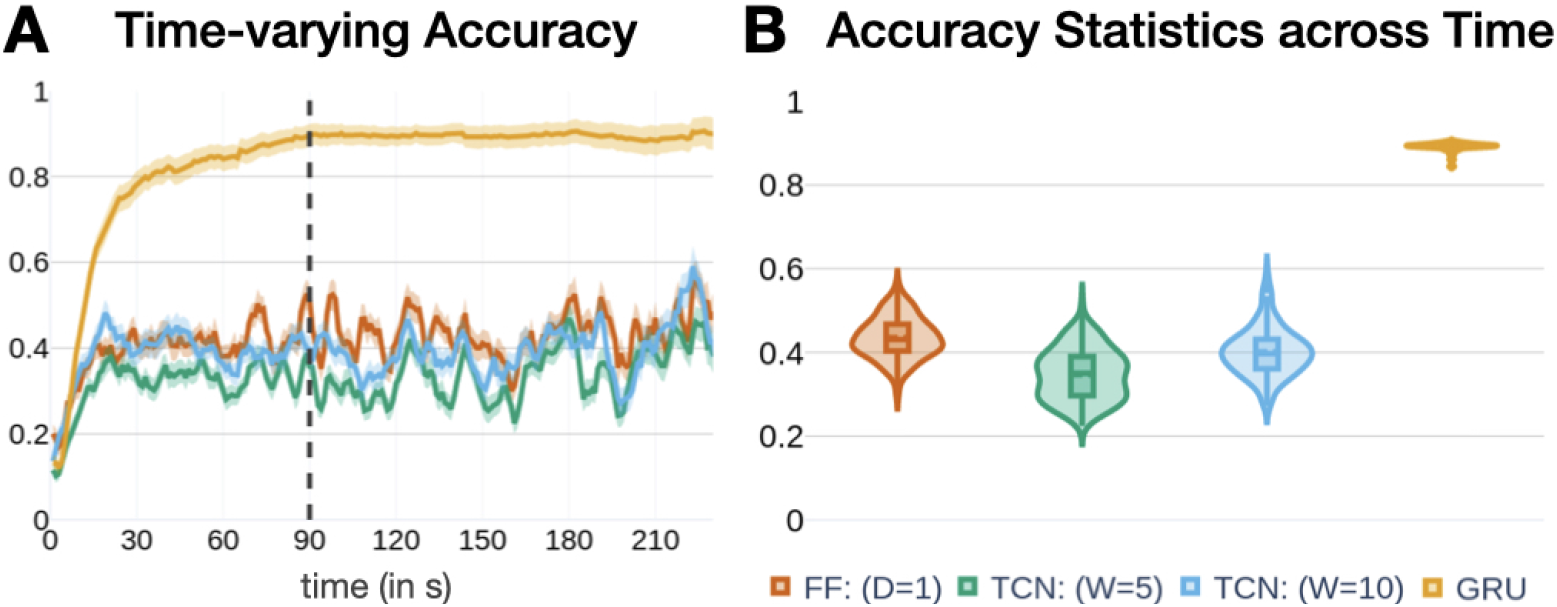
Prediction of movie clip (15-way classification). (A) Average clip prediction accuracy using the neural network architecture with gated recurrent units (GRUs) as a function of time. Accuracy increased sharply during the first 60 seconds, and stabilized around 90 seconds. Results using a feed-forward architecture (FF; 1 layer, 103 units), and temporal convolutional networks (TCN; kernel widths of 5 and 10) also shown. Error bars show the 95% confidence interval of the mean across test participants. (B) Summary of accuracy results after 90 seconds (transient period).

### Dimensionality reduction

Several dimensionality reduction techniques exist that could be used to probe lower-dimensional representations in our system. To capture the temporal relationships in the time series data in the dimensionality reduction process, we combined the GRU network with a simple additional fully connected layer for dimensionality reduction (DR-FC). We refer to this model as the *GRU encoder* (Fig 1B). Formally, GRU outputs, ***h**_t_*, were linearly projected onto a lower-dimensional space containing units with linear activation:

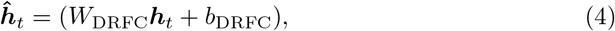

where the weight matrix (*W*_DRFC_ had dimensions *N_h_ × N_ĥ_; N_ĥ_ < N_h_*). We refer to the layer as the *Dimensionality Reduction Fully Connected (DR-FC)* layer. The output of this layer, ***ĥ**_t_*, is sensitive to past inputs, ***x**_t_*, thus effectively capturing temporal dependencies. An additional fully connected layer was used to predict labels, as in the previous subsection.

For dimensionality reduction, only the DRFC and final FC layers were trained (via standard backprogapation). In other words, the training weights of the GRU network for classification were frozen in place after that initial learning phase. For visualization, when the dimensionality of ***ĥ**_t_* was reduced to three, we plotted “trajectories” in a state-space representation (Fig 4A).

**Fig 4.**
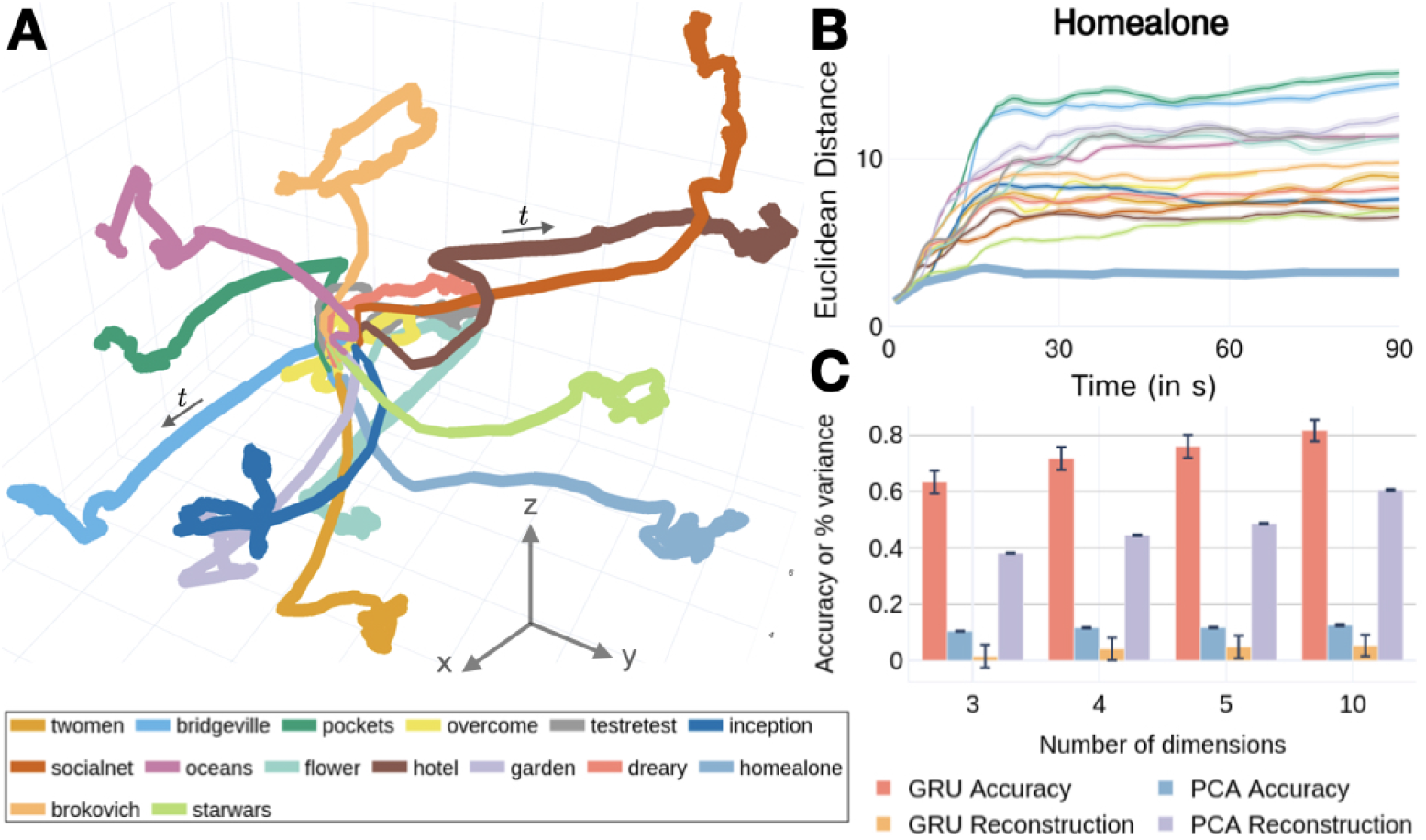
Low-dimensional trajectories. (A) Trajectories for all clips. Solid line: mean trajectory averaged across participants (line thickness is scaled by variance, which was highest at the end of the clip). All trajectories progressed away from the center (see sample arrows). The inset provides clip abbreviations. (B) Euclidean distance between trajectories. The Euclidean distance between the clip trajectory while watching *Home Alone* and the mean trajectory across participants for a second clip was computed. The thicker line corresponds to the distance of participants’ *Home Alone* trajectories to the mean of this clip. The same results are shown for all clips in S1 Fig. (C) Clip prediction accuracy and fraction of variance captured after reconstruction using low-dimensional models. Error bars correspond to the standard error of the mean across participants.

### GRU decoder

Can low-dimensional representations obtained for classification be used to reconstruct the original data? Another GRU-based module was used to reconstruct brain activation from the learned low-dimensional representations, ***ĥ**_t_*, which we refer to as the *GRU decoder* (Fig 1C). The GRU decoder was trained independently (that is, separately from the GRU encoder), by minimizing the mean squared error between the GRU decoder output, 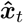, and the original time series input, ***x**_t_*, at each time step.

### Saliency maps for spatiotemporal importance

To help evaluate the contribution of individual inputs (ROIs or voxels) to classification, we employed a *saliency* measure [35]. Saliency assigns importance scores to each input feature based on their contribution to the model’s predictions. Once the model is fully trained, saliency can be computed at every at time, *t*. Given input activation, ***x**_t_*, the GRU classifier generates an output vector, ***ŷ**_t_*. As the process is supervised, the true class label, *c* ∈ {1*, ···, C*}, is known. Thus, the *c*^th^ element of the output vector, 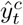, is the class score for the input’s true class label.

Although 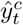 is determined by a complex, non-linear transformation, it can be approximated using a first-order Taylor approximation:

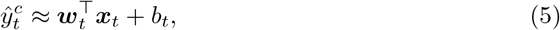

where the operator ⊤ is the transpose, and the the weight vector, ***w**_t_*, is defined as the gradient of 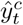 with respect to the activation input:

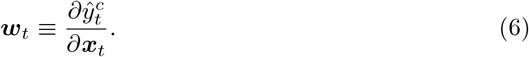

The value of the *i*^th^ element, 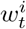, was the saliency value for the *i*^th^ input (ROI or voxel) at time *t*. The saliency captures the contribution of the input activation at time *t* to the true-class score, 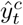. Thus, the higher the value, the stronger the effect on predicting the class in question. To consider saliency values across participants, we z-scored the gradient values, 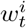, across inputs and time steps for each participant. Because positive or negative contributions were potentially equally important (i.e., increases or decreases in brain activation altered the class prediction), we considered the absolute values of the z-scores above. Finally, we averaged these values across participants to obtain group-level saliency estimates.

To evaluate the robustness of the saliency results, we compared them to those obtained from a null model. To generate a null distribution, using test data, class labels were randomly shuffled, and saliency values computed as defined above. The procedure was repeated 1000 times, yielding a distribution of null-saliency values at each time step (see Fig 5).

**Fig 5.**
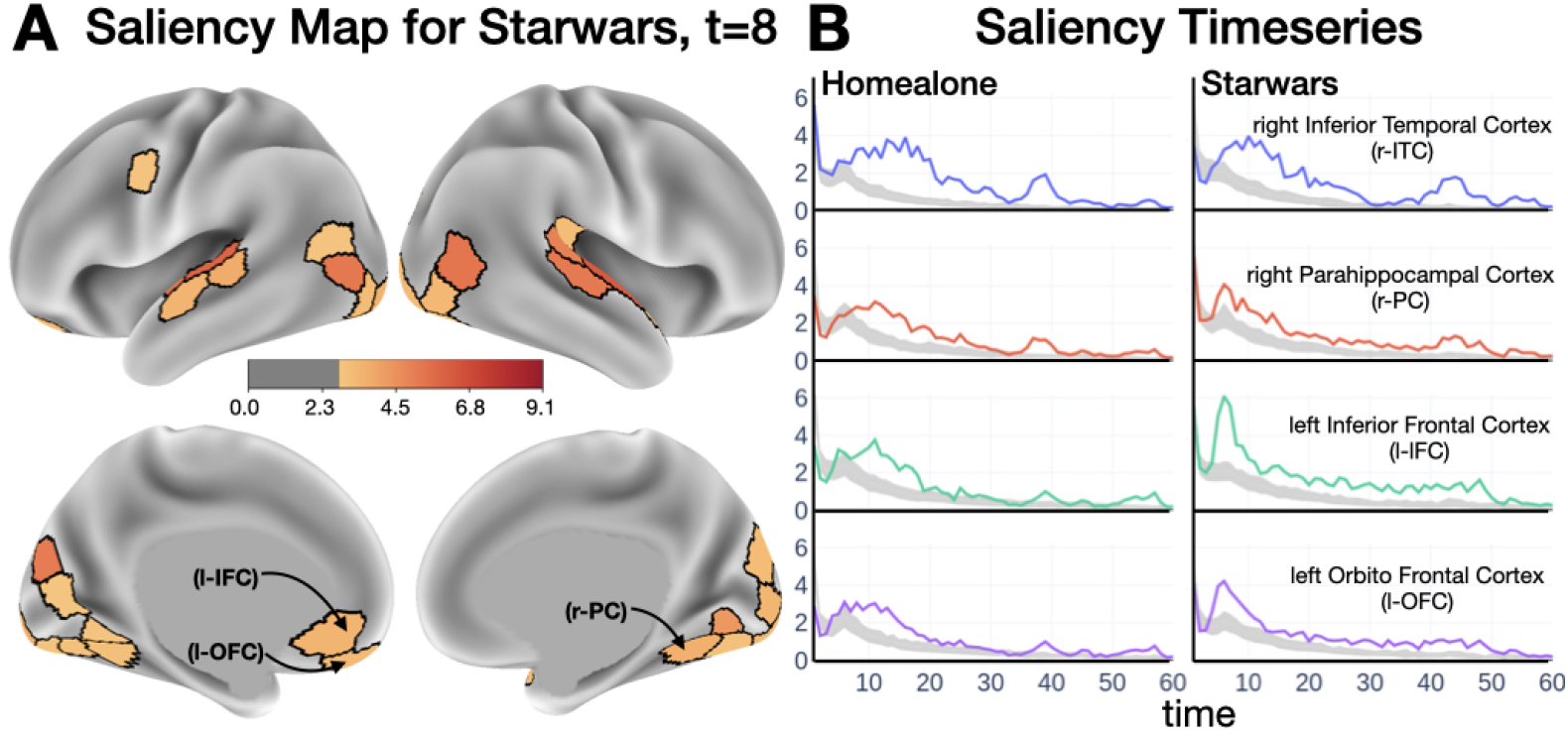
Determining brain contributions to classification. (A) Saliency map for the “Star Wars” clip. For illustration, the top 30 regions are shown. Color scale in arbritary units. See also the video for dynamics. (B) Saliency values for the first 60 seconds of two clips. The gray bands corresponds to the 95-th percentile of null saliency values generated via permutation testing.

### Baseline models

We compared the GRU classifier to two baseline models. Training was based on computing cross-entropy loss between predicted and true labels at each time step (similar to Eq 3). The standard backpropagation algorithm was employed (Adam optimizer), as implemented in TensorFlow.

#### Feed-forward network

The feed-forward (*FF classifier*) network consisted of three fully-connected (FC) layers. Brain inputs at time, ***x**_t_*, were fed into a fully-connected hidden layer, and then transformed into a vector of class scores, as with the GRU classifier. Formally, the hidden layer, 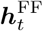, and output layer, 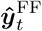, activations were obtained as follows:

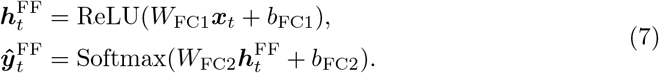

where the ReLU and Softmax operations are standard rectified linear and generalized logistic functions, respectively. The vectors 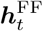 and 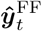 had dimensions 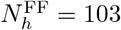 and 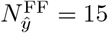 (for 15-way classification) respectively. The number of hidden units was set to 103 in order to maintain the total number of parameters of the classifier (32563) comparable to that of the GRU classifier (32559).

#### Temporal Convolutional Network

We investigated whether static fixed-sized windows would be sufficient for modeling dynamics. A temporal convolutional network (*TCN classifier*) uses fixed-sized convolutional filters, where the output at a time depends upon a fixed temporal window, which is determined by the filter length *l* [36, 37]. Activations after temporal convolution were then transformed to obtain class scores. Formally, brain inputs at each time, ***x**_t_*, were convolved temporally with *k* filters, each with length of *l* time steps. A single filter of size *N_x_ × l* convolved along the time dimension (input time series) and generated a scalar time series. Therefore, *k* such filters produced a *k*-dimensional convolutional layer activation at each time, 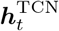, which was used to generate class scores:

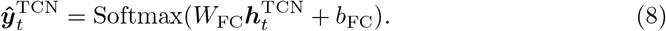

The vectors 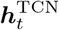 and 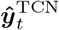 were of dimensions *k* = 25 and 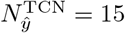, respectively. The choice of *k* = 25 filters, each of width of *l* = 5 time steps, was chosen to keep the the total number of parameters (37915) similar to that of the GRU classifier (32559).

### Predicting behavior

To predict participants’ scores, we employed a GRU-based model, essentially a *GRU regressor*. Separate models were trained to predict fluid intelligence and verbal IQ [38]. In each case, all movie clips were used for training (rather than training a separate model for each clip) to promote learning representations that were not idiosyncratic to a particular clip, and thus generalizable across clips.

We applied the GRU classifier approach (Fig 1). The GRU outputs, ***h**_t_* (dimensionality: *N_h_* = 32), were input to a fully-connected layer with a single unit having a linear activation function. The models learned to predict a single score, *ŷ_t_*, at every time step *t*:

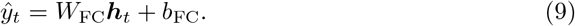

Thus, we were able to generate predicted scores as a function of time, which could be compared with the true score, allowing evaluation of how model predictions evolved. We computed the mean squared error (MSE) between true, *y_t_*, and predicted, *ŷ_t_*, scores at each time point:

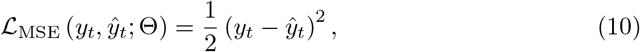

where Θ was the set of learnable parameters. The total loss was defined as the average MSE loss across all time points. The models were trained using backprogagation, using the Adam optmizer for gradient descent.

We compared our approach to connectome-based predictive modeling (CPM; [39]), possibly the state-of-the-art in this regard. In this approach, a functional connectivity matrix is initially formed for each clip based on the Pearson correlation between every pair of brain region time series. Subsequently, the entries in the matrix (or edges) that correlate with the individual differences measure beyond a set threshold (here, 0.2) are retained and a linear model is fit to predict scores based on the functional connectivity values. Given that functional connectivity is employed, CPM predictions rely on activations during the entire clip.

We tested our model statistically at two levels. At each time *t*, once the model was trained (training set only), we evaluated chance performance based upon a null distribution obtained through permutation testing. For each iteration, we randomly shuffled the true scores of test data across participants and then calculated Spearman’s rank correlation between predicted and shuffled scores (1000 iterations). The 95% confidence interval is shown by the gray zone in Fig 6. Because of multiple comparisons across time, we performed an additional test considering all time points simultaneously. The permutation-based test draws from ideas used in the MEG/EEG field to evaluate if two time series differ in time [40]. Consider a sliding window of length *l* over the entire clip time series, and compute the value of the average correlation over the sliding window. This allows us to determine the window that has the highest average correlation. For example, in Fig 6, the window of length *l* = 10 extended from *t* = 139,…, 148, for fluid intelligence. To determine the probability that such value would be observed by chance, we computed the same highest average correlation based on values for shuffled participants (1000 iterations), providing a null distribution (Fig 6, supplemental figures).

**Fig 6.**
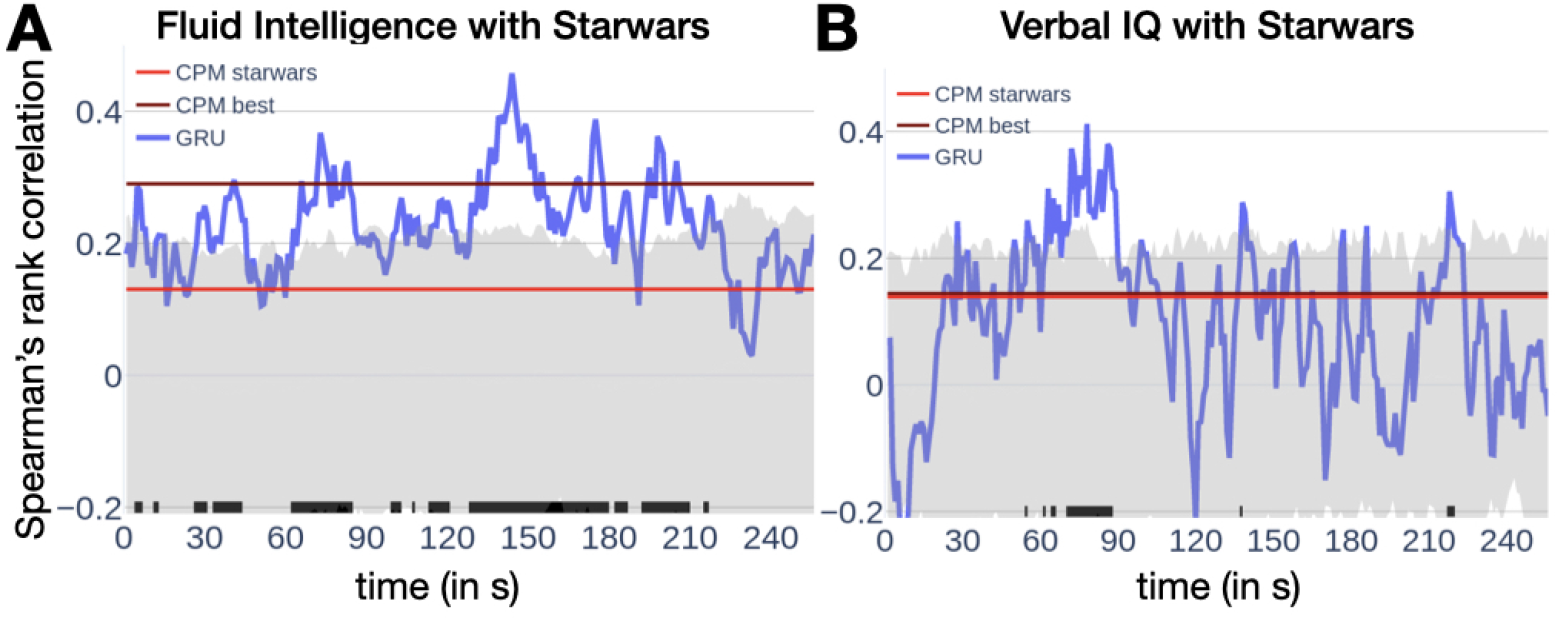
Prediction of behavior. (A) Prediction of fluid intelligence scores as a function of time. Prediction (blue) fluctuates considerably but consistently exceeds chance values (indicated by the tic marks at the bottom). Values obtained by connectome-predictive modeling (CPM) are indicated for comparison (red: CPM applied to *Star Wars* data; maroon: highest value applying CPM across all clips). The gray region indicates the 95^th^ percentile region based on permutation testing (N.B.: applies to our method only, not CPM). (B) Prediction of verbal IQ as a function of time. Only short periods of time of the Star Wars clip exceeded chance levels.

### Summary of models used

Table 1 summarizes all model architectures and documents all hyperparameters used for training.

**Table 1.** Summary of models implemented in the paper. Hyperparameters and other parameters: LR: controlled the rate of update for gradient descent; Dr: dropout for the GRU layer [41]; L2: L2 regularization coefficient for the GRU layer; Batch size: number of training samples passed through the model before updating model parameters; Epochs: number of passes through the training dataset during training.

| Model<br>(Dataset) | Architecture<br>Layer (unit(s)) | LR | Dr | L2 | Batch size | Epochs |
| --- | --- | --- | --- | --- | --- | --- |
| GRU classifier<br>(movie-watching) | GRU (32 units)<br>FC (15 units) | $6 \times 10^{-3}$ | $10^{-6}$ | $10^{-3}$ | 32 | 45 |
| FF classifier<br>(movie-watching) | FC (103 units)<br>FC (15 units) | $10^{-3}$ | NA | NA | 32 | 45 |
| TCN classifier<br>(movie-watching) | TCN (25 filters)<br>FC (15 units) | $10^{-3}$ | NA | NA | 32 | 50 |
| GRU encoder<br>(movie-watching) | GRU (32 units)<br>FC ($N_h$ units)<br>FC (15 units) | $6 \times 10^{-3}$ | $10^{-6}$ | $10^{-3}$ | 32 | 45 |
| GRU regressor<br>(movie-watching) | GRU (1 unit)<br>FC (1 unit) | $10^{-3}$ | NA | NA | 30 | 20 |
| GRU classifier<br>(moving-circles) | 3×GRU (32 units)<br>FC (1 unit) | $6 \times 10^{-3}$ | $10^{-6}$ | $10^{-3}$ | 32 | 28 |

## Results

Our goal was to develop a spatiotemporal decoding framework to predict a stimulus, mental state, behavior, and/or personality traits based on brain signals. A key element of our model was a recurrent neural network based on Gated Recurrent Units (GRUs; [31]) sensitive to current and past inputs (Fig 1A). To determine the generalizability of our findings, all results reported below were obtained with not used in any fashion for training and/or tuning. For a complete description of the architecture, please refer to Materials and Methods.

### Generalizability of spatiotemporal patterns

To test the system’s ability to generalize spatiotemporal brain patterns across participants, we employed the *GRU classifier* to predict movie clip labels from functional MRI data available from the Human Connectome Project. We performed 15-way classification as a function of time (Fig 3A). Accuracy increased sharply during the first 60 seconds, and stabilized around 90 seconds. Mean classification accuracy after this transient period was 89.46% (Fig. 3B). For completeness, a formal evaluation of chance performance based on the null distribution obtained through permutation testing resulted in a mean accuracy of 8.40% (1000 iterations; [42]).

### Is temporal information necessary for clip prediction?

If capturing temporal information and long-term dependencies are important for classification, performance should be affected by temporal order. To evaluate this, we shuffled the temporal order of fMRI time series for test data. To preserve the autocorrelation structure in fMRI data while shuffling, we used a wavestrapping approach [43]. Classification accuracy reduced considerably to 48.44%.

We then compared the GRU classifier to alternative schemes. Our objective was not to benchmark our proposal against possible competitors, but to further probe properties captured by our architecture. First, using a simple feed-forward network with one hidden layer, classification accuracy was 43.51%. We constrained our choice of units (103) such that the the total number of parameters of the feed-forward classifier (32, 563 parameters) was approximately equal to that of the GRU classifier (32, 559 parameters). As feed-forward classifiers do not capture temporal dependencies, we next employed temporal convolutional networks. We used 25 kernels with a width (*W*) of 5 time steps such that the number of parameters of the classifier (37, 915 parameters) approximated that of the GRU classifier (see Methods for kernel details). Classification accuracy was only 35.65%. We then increased the kernel width to 10 time steps (75, 415 parameters). This numerically improved accuracy to 41.85%.

Together, the results above are consistent with the notion that temporal information is essential for high-accuracy classification, and that our GRU-based architecture is capable of capturing long-term dependencies that are missed by temporal convolutional networks, for example. Note, however, that temporal shuffling of the time series still allowed the GRU classifier to attain classification at around 50% correct. As the temporal shuffling was unique for each training permutation, order information was broken down, indicating that non-temporal information supported considerably above-chance performance.

### Low-dimensional trajectories as spatiotemporal signatures

We investigated the dimensionality of the inputs required for successful stimulus prediction by using a supervised non-linear dimensionality reduction approach (Fig 1B). The original dimensionality of the input, ***x**_t_*, was the number of regions (*N_x_* = 300). After learning, the GRU outputs provide a low-dimensional latent spatiotemporal representation of the input, which is encoded in the vector ***h**_t_* (*N_h_* = 32). To further reduce dimensions, GRU signals, ***h**_t_*, were linearly projected onto a lower-dimensional space, ***ĥ**_t_*, using a fully connected layer. We refer to this weight matrix as the *Dimensionality Reduction Fully-Connected (DRFC)* layer, and to this model as the *GRU encoder*. Since ***ĥ**_t_* is a low-dimensional representation of the history of ***x**_t_*, the inputs are not treated independently, effectively leading to non-linear temporal dimensionality reduction. As in the original classifier, a final FC layer was used to predict labels based on ***ĥ**_t_*.

Performance with low-dimensional encoding was surprisingly high; 3 dimensions yielded 63.30% accuracy. In Fig 4A, we show low-dimensional signals for each clip, which we refer to as *trajectories*: consecutive low-dimensional ***ĥ**_t_* states, (*ĥ*_1_(*t*)*, ĥ*_2_(*t*)*, ĥ_3_(t*)), describe the trajectory in *state space*. In other words, at each time *t*, the value of each unit is plotted along the (*x, y, z*) axes. To quantify a notion of proximity between trajectories, we computed the Euclidean distance between them as a function of time (Fig 4B). To compute the distance between clips *A* and a reference clip *B*, we first computed the mean trajectory of *B* averaged across participants. Then, for each participant’s clip *A* trajectory, we computed the Euclidean distance between *A* and the mean trajectory of *B* at every time step. Accordingly, the proximity of a clip with itself was not zero (indicated by a thicker line), and reflected the variability of participant trajectories around the clip’s average. As expected, the evolution of low-dimensional trajectories closely reflected the temporal accuracy obtained using the original GRU classifier. Clip trajectories were initially close to one another, but slowly separated during the first 60-90 seconds of the clip.

We further investigated low-dimensional projections with 4, 5, and 10 dimensions, at which point performance (81.56%) was not far (within 10%) from that of full dimensionality (Fig 4C). These results reveal that latent representations with as few as 10 dimensions captured essential discriminative information. The effectiveness of the GRU encoder in capturing low-dimensional information can be appreciated by applying principal components analysis on the input data, ***x**_t_*, which yielded very low prediction accuracy, although it was capable of recovering 60% of the input signal variance with 10 dimensions (Fig 4C).

To investigate the content of the latent space uncovered by the GRU encoder, we performed the following analysis. By construction, the low-dimensional vector captures information to classify the input signals. How much information about the input does this projection preserve? To evaluate this, we considered the low-dimensional signal, ***ĥ**_t_*, and used it to try to reconstruct the input signal, ***x**_t_*. Within our framework, it is natural to do so using a GRU decoder (Fig 1C), which reconstructs the input time series (the reconstructed signal is called 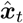). Doing so captured a very modest amount of signal variance (less than 10%).

### Spatiotemporal importance maps

The GRU representation that supports classification lies in a latent space that is fairly disconnected from the original brain activation signals, ***x**_t_*. It is therefore important to relate GRU states to brain activation. We defined a *saliency* measure to capture the contribution of a brain region to classification as a function of time (see Methods). Fig 5A shows a saliency map at *t* = 8 for the *Star Wars* clip, also shown dynamically for the first 60 seconds (S2 Video). The saliency time series in a few brain regions shows that values were relatively high initially and gradually decreased over the first minute of the clip (Fig 5B). The evolution of saliency for the *Home Alone* clip was somewhat similar (see also the saliency map video for the *Brokovich* clip in S1 Video).

### Predicting behavior

Recent studies have employed brain data to predict participants’ behavioral capabilities or personality-based measures [44–48]. We tested the extent to which spatiotemporal information captures individual-based measures based on our recurrent neural network framework. Here, we targeted individual scores for fluid intelligence and verbal IQ. The same approach used for classification was employed, with the model predicting a scalar, namely, a participant’s score. We used all clips for training (rather than training a separate model based on each clip) to promote learning representations that are not idiosyncratic to a particular clip.

Fig 6 shows model performance temporally for the *Star Wars* clip, one of the best predicting clips. At every time point, predicted scores are correlated with actual participants’ scores. For fluid intelligence, the observed correlations were often above chance levels (gray zone indicates 95% confidence level at specific time *t*), but more modestly for verbal IQ. As our model made predictions as a function of time, we developed a permutation-based test to evaluate the predictions while controlling for multiple comparisons (fluid intelligence, *p* = 0.0013; verbal IQ: *p* = 0.0038). The supplementary figures (S2 Fig, S3 Fig, S4 Fig, S5 Fig) show results for all clips. For reference, we compared our approach to connectome-based predictive modeling based on patterns of functional connectivity (CPM; [39]), possibly the state-of-the-art in this regard (see also [44, 46]). We include values predicted with this technique to provide an informal comparison, as this method provides a single prediction per clip, unlike out approach which is time varying.

### Dynamic multivariate pattern analysis

Thus far, we applied our model to region-based activation patterns. The approach can also be employed to perform voxel-based (or grayordinate-based) dynamic multivariate pattern analysis (MVPA). Here, we applied our model to the prediction of experimental conditions from a dataset collected in our laboratory [3]. Briefly, in the “moving circles” paradigm, over periods of three-minute blocks, two circles moved on the screen, at times approaching and at times retreating from each other. If they touched, participants received a non-painful but highly unpleasant electrical shock. The movement of the circles was smooth but not predictable. In particular, the circle motions were set up to include multiple instances of “near misses”: periods of approach followed by retreat; in such instances, the circles came close to each other and retreated just before colliding. Based on near-misses, we defined *approach* and *retreat* states: each lasted seven time steps (total of 11.25 seconds) during which the circles approached or retreated from each other.

We performed dynamic MVPA by using voxels from the anterior insula, a region strongly engaged by the experimental paradigm [2, 3]. Classification accuracy ranged from 59-63% over the approach and retreat segments (the upper bound of the 95% confidence interval determined with permutation testing was 50.37%). We also computed saliency in a voxelwise manner; values were substantially higher at the outset of a approach/retreat segment and relatively lower for the subsequent time points. Fig 7 shows a snapshot at *t* = 6 seconds, illustrating the spatial organization of the map.

**Fig 7.**
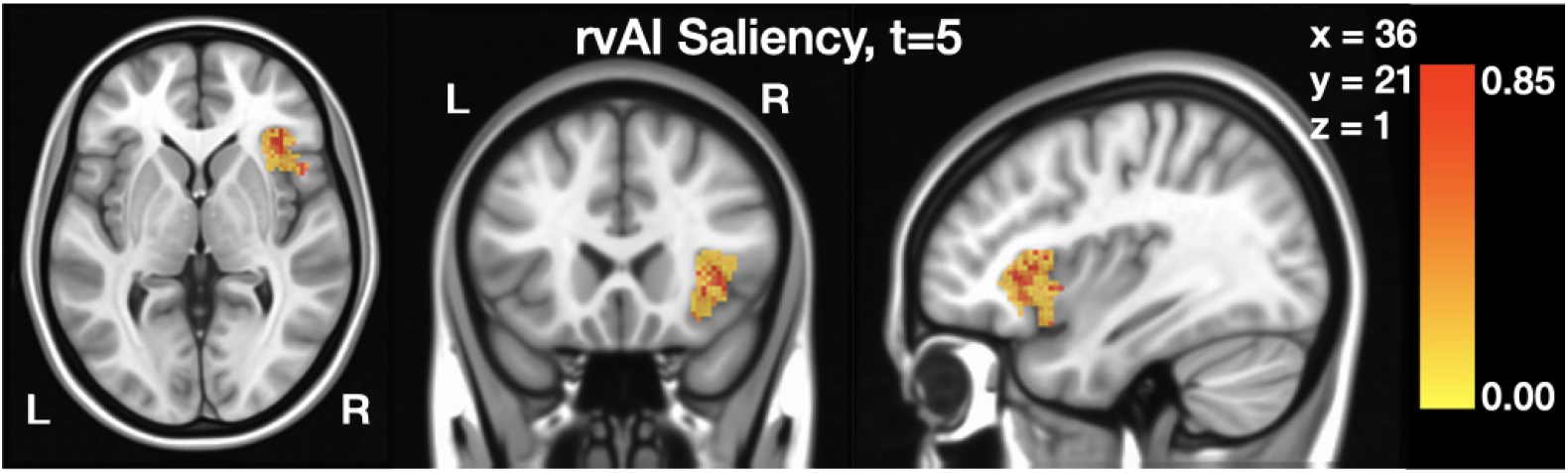
Voxelwise prediction of experimental conditions: threat vs. safe. Only voxels from the anterior insula were employed. Saliency values at *t* = 6 are shown; for illustration no thresholding was applied.

## Discussion

We sought to characterize distributed spatiotemporal patterns of functional MRI data during movie watching and continuous task conditions. To do so, we employed recurrent neural networks sensitive to the long-term history of the input signals, and applied them to the problem of input classification. The model was tested on brain data both at the level of region of interest and voxels. All neural network training and model tuning were performed using a training set independent of the test set to establish the model’s ability to generalize learned representations to unseen participants. The framework we developed is general and its subcomponents can be easily exchanged to utilize other algorithms (e.g., long short-term memory networks, or novel developments).

### Spatiotemporal representations

Spatiotemporal information learned in a given set of participants generalized very well to new participants, where challenging 15-way classification was ~90% correct for movie clip data. These results are noteworthy because they suggest that activation patterns distributed across space and time are shared across participants when viewing naturalistic stimuli. To understand the type of information captured by our model, we compared it to a few simple schemes, including a simple feedforward network, and temporal convolutional networks. The low performance with these models (~40%) suggests that long-term dependencies are essential for input classification. Indeed, temporal shuffling of the input time series decreased performance to ~50%. As temporal shuffling preserves the input signals but scrambles their sequence, these results reveal that the precise temporal order is key for the ability to generalize to unseen data.

### Latent dimensionality

The question of the “inherent” dimensionality of brain signals has attracted considerable attention in recent years [18–20, 49]. Here, we investigated this issue for the movie clip data, where inputs were 300-dimensional based on the ROI time series. As the number of units in the hidden layer was 32, a considerable amount of dimensionality reduction was accomplished at the outset. Further investigation revealed that this signal could be further reduced to ~10 dimensions without compromising prediction accuracy appreciably. Strikingly, even with three dimensions, accuracy was relatively high (77.30%). Importantly, whereas the mapping onto a lower dimensionality space preserved classification information about the input category, it was very poor at reconstructing the input time series itself. Thus, the content of the latent low-dimensional representation was substantially distinct from the fMRI signal itself.

### Saliency

Although prediction is valuable in applied settings (e.g., clinical diagnosis), our goal was to be able to characterize distributed spatiotemporal representations in the brain, too. A measure of saliency uncovered brain regions and their time-varying importance to movie clip prediction. The contribution of brain areas was highest during the first minute in a manner that paralleled the time course of accuracy. Spatiotemporal saliency maps revealed that, whereas early visual and auditory regions contributed to prediction, multiple additional areas contributed too, supporting the idea that classification was not simply based on low-level movie features. For example, one of the regions with high salience (right inferior temporal cortex) included part of the fusiform gyrus, which is known to be highly responsive to faces. Other parts of inferior temporal cortex, including parahippocampal cortex which is involved in high-level object recognition, participated too. As further examples, the inferior frontal cortex and the orbitofrontal cortex supported clip classification, too. Together, these results support the idea that the overall model captured biologically relevant information, and shows how signal importance fluctuates spatiotemporally.

### Individual differences

Recent work has sought to predict individual differences based on functional MRI data [39, 44–48]. Our model was capable of predicting fluid intelligence and verbal IQ from unseen participants at levels comparable to, and possibly better than, established methods such as connectome-based predictive modeling, which is based on information from functional connectivity matrices [39]. Some movies (e.g., *Star Wars*, *Social Net*, *Oceans*) performed much better at prediction than others, and at particular segments of the movie. This raises the intriguing possibility that those particular segments are better “tuned” to particular individual differences (in this case, fluid intelligence). Of course, larger datasets will be needed to adequately assess this possibility.

### Dynamic multivariate pattern analysis

We applied the model developed here with inputs at the level of voxels, so as to implement dynamic MVPA. We tested the approach in a dataset involving continuous changes in threat level based on the proximity of two moving circles. Voxels from the anterior insula were used to predict stimulus condition (threat vs. safe). Although accuracy was relatively modest (~60%), the results demonstrate the feasibility of the approach. Saliency maps also uncovered subsectors of the anterior insula with increased importance for prediction, illustrating how it is possible to identify voxels that contribute to decoding the experimental conditions, opening an avenue for the application of representational similarity analysis [24] in a dynamic setting.

### Other approaches

Decoding approaches using some temporal information have been used in the past [11–16]. Recurrent neural networks have also been employed for brain decoding more recently, albeit in some cases on relatively static task paradigms, such as working memory which generates stable brain states at the temporal scale of fMRI [17, 50]; working memory conditions can be predicted even when temporal information is eliminated [51]. Because most fMRI datasets are typically small, they are often evaluated using cross-validation without assessing their generalization on held-out data (see discussion by [52]). The size of the datasets studied allowed us to test the model in a separate set of participants thereby evaluating the model’s ability to generalize beyond trained data (see also [50]). Finally, we stress that the goal of the present study was not to describe a model that outcompetes others but to describe a general, modular modeling approach to investigating spatiotemporal dynamics of brain data, here applied to fMRI.

## Conclusion

In the present paper, we developed a computational approach to study and characterize spatiotemporal brain signals in the context of fMRI. The framework can be applied to other types of brain data for discovering and interpreting brain dynamics.

## Supporting information

**S1 Fig. Euclidean distances between trajectories for all movie clips.**

Euclidean distances between trajectories. In the inset, the duration of every clip is indicated in parenthesis.

**S2 Fig. Fluid Intelligence predictions for all movie clips.** Fluid Intelligence predictions for all movie clips. Conventions as in the figure in the main text.

**S3 Fig. Fluid Intelligence predictions: null distributions.** Null distributions of fluid intelligence predictions. Vertical blue lines indicate the prediction based on actual data.

**S4 Fig. Verbal IQ predictions for all movie clips.** Verbal IQ predictions for all movie clips. Conventions as in the figure in the main text.

**S5 Fig. Verbal IQ predictions: null distributions.** Verbal IQ predictions: null distributions. Vertical blue lines indicate the prediction based on actual data.

**S1 Video Saliency movie for Brokovich clip.** [[link](https://drive.google.com/file/d/1Why34mgp4wedzLODZUl1Wyv3lbuPaOPN/view?usp=sharing)]

(https://drive.google.com/file/d/1Why34mgp4wedzLODZUl1Wyv3lbuPaOPN)

**S2 Video Saliency movie for Star Wars clip.**[[link](https://drive.google.com/file/d/1jaTgnHPhvouKM28-r3rGohZNPP3lvGfi/view?usp=sharing)]

(https://drive.google.com/file/d/1jaTgnHPhvouKM28-r3rGohZNPP3lvGfi)

**S1 Appendix. Gated recurrent units.**

## Acknowledgments

M.V. was supported by a fellowship by the Brain and Behavior Initiative, University of Maryland, College Park. L.P. is supported by the National Institute of Mental Health (R01 MH071589 and R01 MH112517). Data were provided in part by the Human Connectome Project, WU-Minn Consortium (Principal Investigators: David Van Essen and Kamil Ugurbil; 1U54MH091657) funded by the 16 NIH Institutes and Centers that support the NIH Blueprint for Neuroscience Research; and by the McDonnell Center for Systems Neuroscience at Washington University.

## Footnotes

1 The source code used to produce the results and analyses presented in this manuscript are available from GitHub repository: https://github.com/LCE-UMD/GRU.

