## Supplemental material for "Learning brain dynamics for decoding and predicting individual differences"

April 5, 2021

**S1 Videos; Saliency movies**

- **Saliency movie for Brokovich clip.**[link]
  (https://drive.google.com/file/d/1Why34mgp4wedzLODZUl1Wyv3lbuPaOPN)
- **Saliency movie for Star Wars clip.** [link]
  ( https://drive.google.com/file/d/1jaTgnHPhvouKM28-r3rGohZNPP3lvGfi)

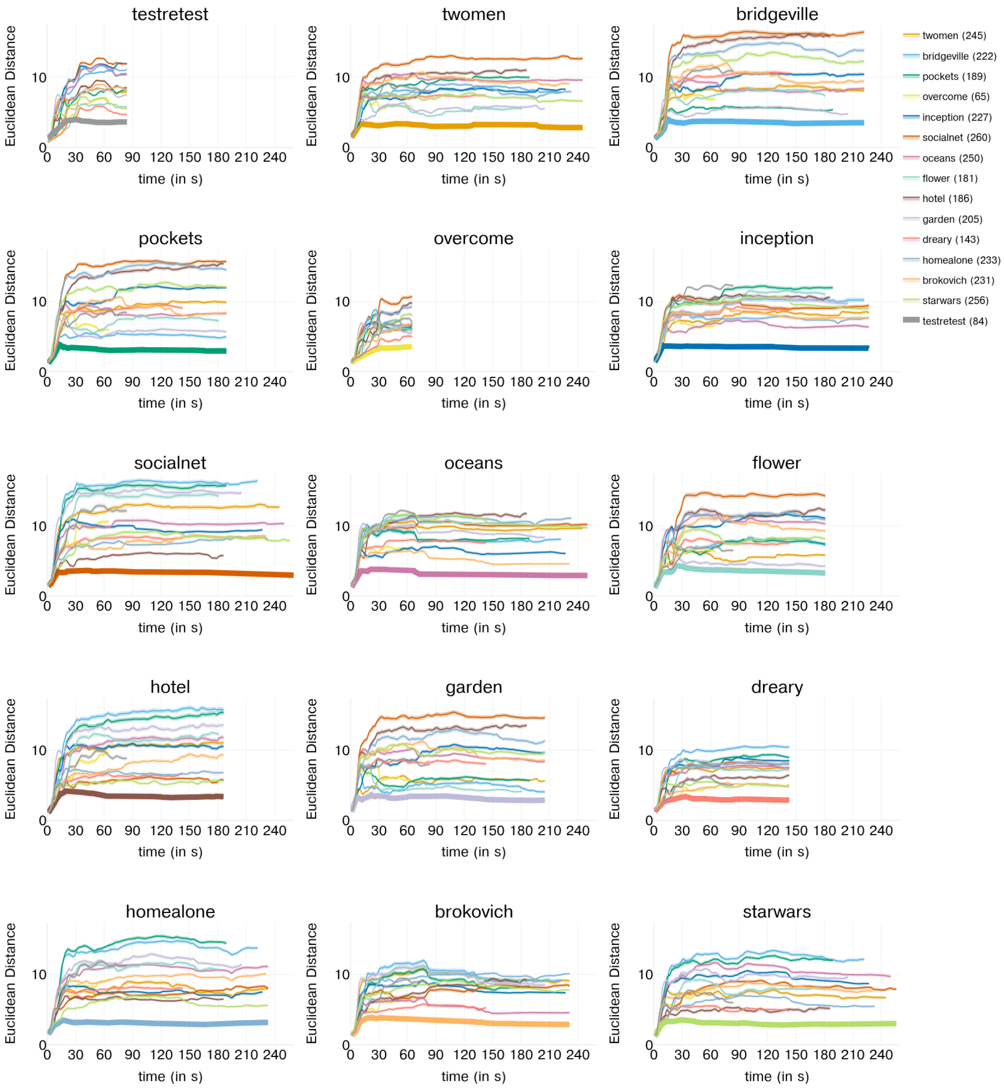


**Fig S1.** Euclidean distances between trajectories for all movie clips. Euclidean distances between trajectories. In the inset, the duration of every clip is indicated in parenthesis.

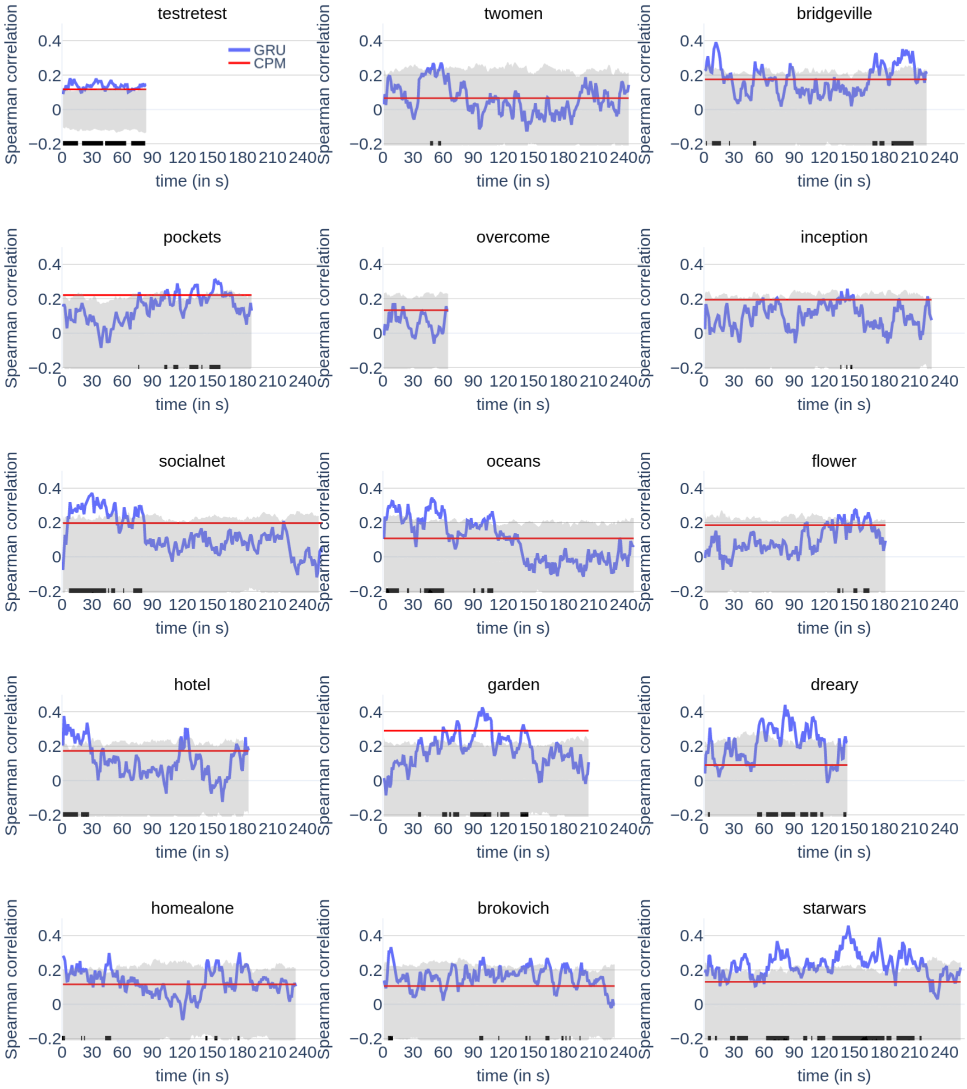


**Fig S2.** Fluid Intelligence predictions for all movie clips. Fluid Intelligence predictions for all movie clips. Conventions as in the figure in the main text.

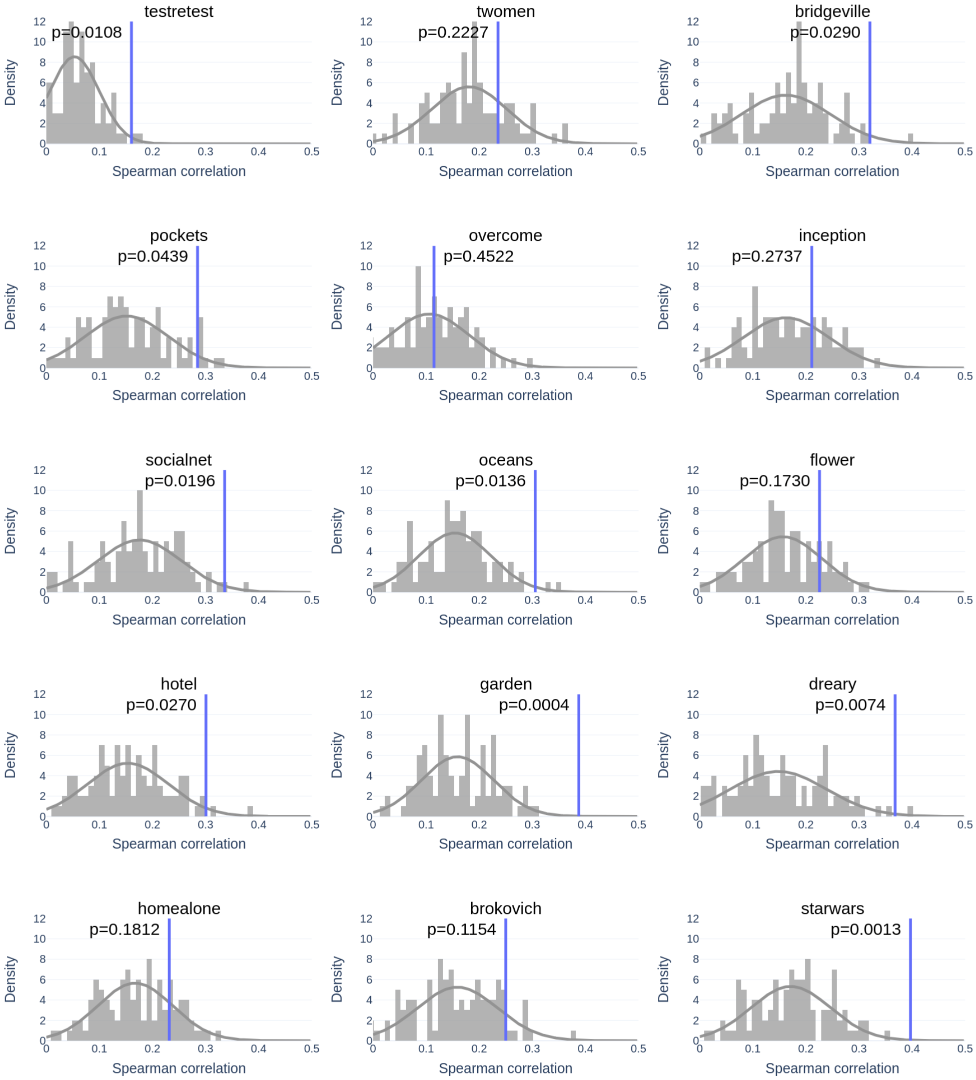


**Fig S3.** Fluid Intelligence predictions: null distributions. Null distributions of fluid intelligence predictions. Vertical blue lines indicate the prediction based on actual data.

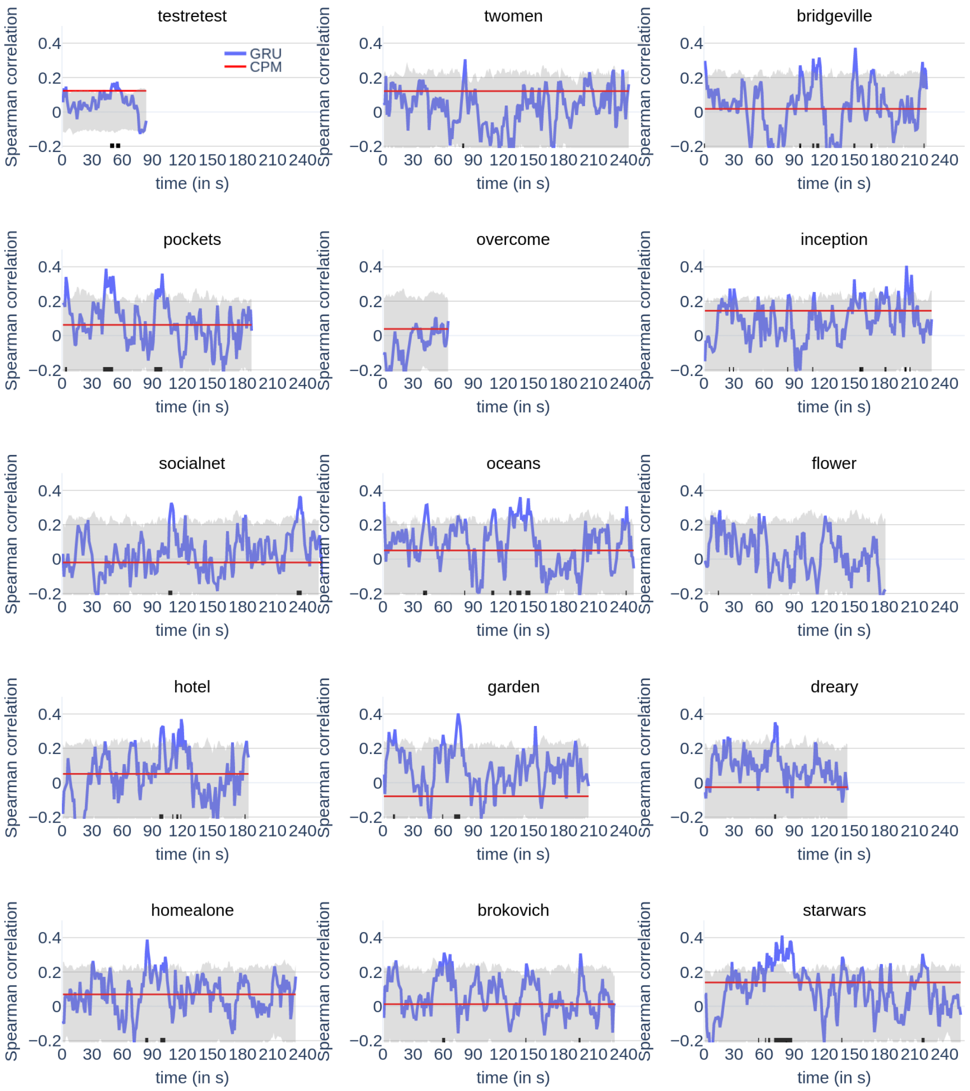


**Fig S4.** Verbal IQ predictions for all movie clips. Verbal IQ predictions for all movie clips. Conventions as in the figure in the main text.

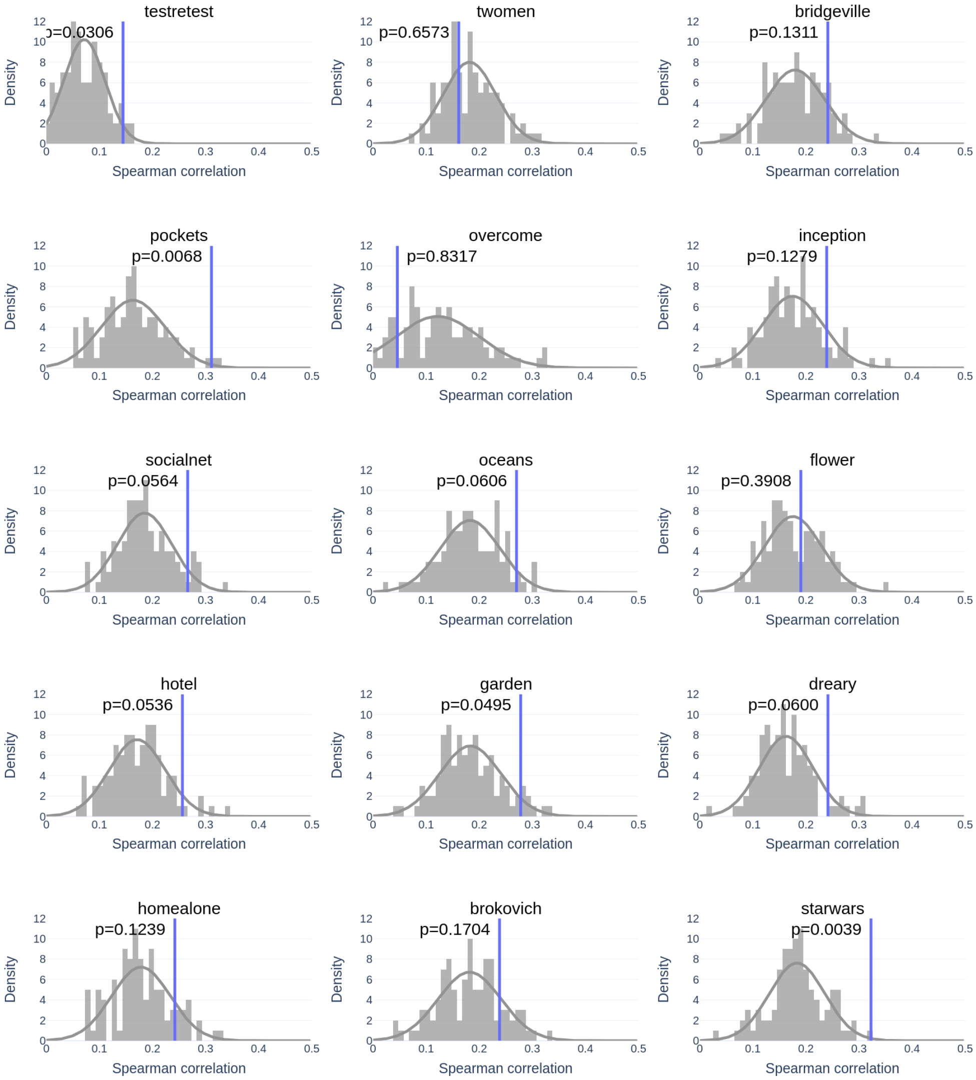


**Fig S5.** Verbal IQ predictions: null distributions. Verbal IQ predictions: null distributions. Vertical blue lines indicate the prediction based on actual data.

**S1 Appendix; Gated Recurrent Units** We employed recurrent neural networks with Gated Recurrent Units (GRUs) which overcome challenges in learning temporal information 1) by adaptively updating temporal history, and 2) by resetting/dropping temporal history that is irrelevant for future predictions [1, 2].

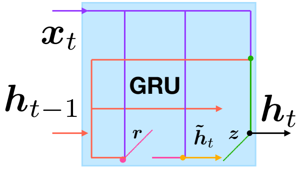


**Fig S6.** Formal specification of a GRU unit. The input, $\boldsymbol{x}_t$, and previous GRU output, $\boldsymbol{h}_{t-1}$, together generate a reset gate vector, $\boldsymbol{r}$, and an update gate vector, $\boldsymbol{z}$.

Given the input, $\boldsymbol{x}_t$, and previous output vector, $\boldsymbol{h}_{t-1}$, GRU units compute a reset gate vector, $\boldsymbol{r}$, and an update gate vector, $\boldsymbol{z}$:

$$\begin{aligned} \boldsymbol{r} &= \sigma\left(W_r\boldsymbol{x}_t + U_r\boldsymbol{h}_{t-1} + b_r\right), \\ \boldsymbol{z} &= \sigma\left(W_z\boldsymbol{x}_t + U_z\boldsymbol{h}_{t-1} + b_z\right), \end{aligned} \tag{1}$$

where $W_r, U_r, W_z, U_z$ are weight matrices and $b_r, b_z$ are bias vectors that are defined during the learning phase. The symbols $\sigma(\cdot)$, $\tanh(\cdot)$, and $\odot$ represent the sigmoid activation function, the hyperbolic tangent function, and the Hadamard product, respectively.

The two gate vectors are then used to compute a candidate state vector:

$$\tilde{h}_t = \tanh\left(W_h\boldsymbol{x}_t + U_h\left(\boldsymbol{r}\odot\boldsymbol{h}_{t-1}\right) + b_h\right), \tag{2}$$

using weight matrices $W_h, U_h$ and bias vector $b_h$, which are defined during the learning phase ($N_{\tilde{h}} = 32$ dimensions). The reset vector, $\boldsymbol{r}$, controls the amount of information from the past output, $\boldsymbol{h}_{t-1}$, used to compute the candidate activation. Thus, when $\boldsymbol{r}$ was close to zero, the prior memory, $\boldsymbol{h}_{t-1}$, was de-emphasized and $\tilde{\boldsymbol{h}}_t$ was "reset" with information from the current (brain signal) input, $\boldsymbol{x}_t$. Finally, the update gate vector, $\boldsymbol{z}$, governs the fraction of the previous state activation to be carried forward:

$$\boldsymbol{h}_t = (\boldsymbol{1}-\boldsymbol{z})\odot\boldsymbol{h}_{t-1} + \boldsymbol{z}\odot\tilde{\boldsymbol{h}}_t. \tag{3}$$

Each GRU unit has its own reset and update gate units, thus allowing each unit to learn dependencies over multiple time scales. Units that learn long-term dependencies have their update gates frequently active, while those that learn shorter-term dependencies have their reset gates frequently active.
